# Find me if you can: Pre- and Post-hurricane Densities of the Red-bellied Racer (*Alsophis rufiventris*) on St. Eustatius, and a review of the genus in the Caribbean

**DOI:** 10.1101/2021.07.05.451169

**Authors:** Hannah Madden, Denny S. Fernández, Raymond L. Tremblay, Kevin Verdel, Brent Kaboord

## Abstract

We estimated population densities of the red-bellied racer (*Alsophis rufiventris*) on the Caribbean island of St. Eustatius in 2011, 2018 and 2019 to determine the likely influence of hurricanes Irma and Maria (September 2017), in addition to evaluating abiotic parameters which may be correlated with its presence. Surveys were conducted at seven sites in 2011 prior to the hurricanes, and at 81 and 108 sites in 2018 and 2019 respectively posterior to the hurricanes. A total of 8.2 ha was surveyed in 2011, and 11.42 ha in 2018/2019. The pre-hurricane (2011) racer density estimate was 9.2/ha (min 7.3 - max 11.6); post-hurricane estimates were 4.6/ha (min 3.4 - max 6.0) in 2018 and 5.0/ha (min 3.8 - max 6.5) in 2019. The pre-hurricane encounter rate of individual racers was 16.0 snakes/hour compared to 0.34 snakes/hour in 2018 and 0.41 snakes/hour in 2019 (post-hurricane). The decrease in encounter rates between 2011 and 2019 implies a negative impact of the hurricanes on racer abundance. Based on calculations of detection probability (0.02 in 2018 and 0.03 in 2019), post-hurricane lambda estimates were 1.82 (95% CI 0.66 - 5.01) in 2018 and 1.60 (95% CI 0.39 - 6.65) snakes/ha in 2019. Given the current small size of the remaining population and the presence of invasive species across the snake’s range, this species could be at risk of local extirpation. We suggest conservation actions such as invasive species management and habitat restoration to enable further recovery.

## INTRODUCTION

The frequency of hurricanes in some tropical areas can be a strong determinant of the structure and composition of biotic communities (Odum, 1970; Walker et al., 1991; Wunderle et al., 2004). Although structural vegetation damage is a primary direct effect of severe hurricanes to mature forests (Boose et al., 2004; Eppinga and Pucko, 2018), food limitation is a significant indirect effect (Cely, 1991). Animals may respond to these indirect effects by altering their diet or relocating to less affected areas or habitats (Wundele et al., 2004). The Lesser Antilles is a group of islands in the Caribbean archipelago, an area characterized by frequent hurricanes, with multiple or major hurricanes striking every eight to nine years (Elsner et al., 1999; Tartaglione et al., 2003). In Puerto Rico, Hurricanes Hugo (1989) and Georges (1998) brought storm force winds that defoliated trees, resulting in reduced cover for the Puerto Rican boa (*Epicrates inornatus* Reinhardt, 1843) and limited access to arboreal sites (Wunderle et al., 2004). As a result, boas were more visible in the post-hurricane (0.20% ± SE 0.03) compared with the pre-hurricane (0.11% ± 0.02) period, suggesting that the species was more abundant than previously thought (Wunderle et al., 2004). Nevertheless, the forest systems of these islands have adapted to withstand frequent storms (Odum, 1970; Doyle, 1981), and species occurring within these systems may have broad habitat breadths (Waide and Reagan, 1996), including snakes in the genus *Alsophis*.

Habitat selection is a fundamental determinant of animal distribution and densities. Reptiles and amphibians often tend to have their distribution restricted by environmental variables. Example of this are well known, and variation in environmental variables that influences reptiles’ distribution include canopy cover (Wunderle et al., 2004; Schlaepfer and Gavin, 2001), substrate (Wunderle et al., 2004), humidity (Brown and Shine, 2002), temperature (Brown and Shine, 2002), vegetation type (Stumpel, 2004), time of day (Stumpel, 2004), as well as hurricane impacted sites (Wunderle et al., 2004). An example of environmental variation influencing snake behavior includes forest cover; this was observed in the black rat snake (*Pantherophis obsoletus* Say, 1823) which thermoregulates at forest edges (Blouin-Demers and Weatherhead, 2001), while the Santa Catalina rattlesnake (*Crotalus catalinensis* Linnaeus, 1758) in the Gulf of California avoids high soil temperatures in specific periods of the year (Martins et al., 2008). Preferred structured canopy and microclimatic variations influence habitat selection in the broad-headed snake (*Hoplocephalus bungaroides* Schlegel, 1837) in Australia (Pringle et al., 2003), while Malayan pit vipers’ (*Calloselasma rhodostoma* Kuhl, 1824) movements were positively correlated with relative humidity (Daltry et al., 1998). In small Colubrids, preferred locations have been shown to be influenced by soil moisture conditions. (Elick et al., 1972).

The Caribbean is classified as a biodiversity hotspot due to its high levels of species endemism (Mittermeier et al., 2004). Unfortunately, the restricted geographic ranges of endemic species puts them at greater risk of extinction from local impacts (e.g., habitat loss, invasive species; Catford et al., 2012), especially specialist species with limited dispersal abilities, restricted population sizes and reduced adaptive capacity (Chichorro et al., 2019; Staude et al., 2020). Snakes in the genus *Alsophis* once occupied over 100 islands, from the Bahamas to Dominica in the Lesser Antilles (Henderson and Sajdak, 1996). However, habitat degradation, human persecution, and the introduction of the Indian mongoose (*Herpestes javanicus* Geoffroy Saint-Hilaire, 1818) and black rat (*Rattus rattus* Linnaeus, 1758) have resulted in more extirpations and extinctions than any other reptilian or amphibian genus in the region (Henderson and Tolson, 2006). Today, several Lesser Antillean species, including *Alsophis antiguae* (Parker, 1933), *Alsophis rijgersmaei* (Cope, 1869), and *Alsophis rufiventris* (Duméril, Bibron and Duméril, 1854), are considered threatened or endangered (Daltry et al., 2001; Henderson and Powell, 2009). Five species of *Alsophis* exist in the Lesser Antilles, each occurring on a single or small number of islands. *Alsophis antillensis* (Schlegel, 1837) has the largest range, occurring on the Monserrat, Guadeloupe, and Dominica island banks. *Alsophis rijgersmaei* occurs only on the main islands of the Anguilla Bank, *A. antiguae* on the Antigua Bank, *A. sanctonum* (Barbour, 1915) on the Isles des Saintes Bank near Guadeloupe, and *A. rufiventris* on the Saba and St. Christopher Bank (Maley, 2004). We have been unsuccessful at locating information on population size estimates or densities for these species, except for general comments highlighted below:

### Historical and Present Knowledge of *Alsophis* Species

The Antiguan racer (*Alsophis antiguae*) has not been collected in at least 80 years and is likely extinct due to mongoose predation (Sajdak and Henderson, 1991). However, Great Bird Island off the Antiguan coast supports a population of *A. antiguae sajdaki* (Henderson, 1990), which in 2001 was thought to total approximately 80 individuals (Daltry et al., 2001). It is unlikely that Great Bird Island (0.083 km^2^) can support more than approximately 100 individuals because of resource limitations (Daltry et al., 2001). Following translocations from Great Bird Island to Rabbi Island, Green Island (2002) and York Island (2008), the total area of occupancy for *A. antiguae sajdaki* increased to 63 ha. Since translocation, the species appeared to thrive in its new habitats where it immediately began reproducing. Since then, the Antiguan snake metapopulation is estimated to have increased to > 1,100 individuals (Daltry et al., 2017). Nevertheless, these racers and their habitats remain potentially vulnerable to invasive mammals such as rats (*Rattus* spp.), cats (*Felis catus* Linnaeus, 1758), dogs (*Canis lupus* Linnaeus, 1758) and mongooses, in addition to harmful developments, and hurricanes (Daltry et al., 2001; 2017).

*Alsophis antillensis antillensis* was thought to be extinct from Guadeloupe, where both Basse-Terre and Grande-Terre have mongoose populations. In 1990, Sadjak and Henderson (1991) visited Terre-de-Haut and detected eight racers in three days. They also visited Terre-de-Bas, where they detected six *A. a danforthi* in three days. More recently, Breuil (2009) reported sightings from a survey in 2003 of *Alsophis antillensis* in Basse-Terre and Grande-Terre and on Ilet à Cabrit (new locality for Les Saintes as *Alsophis sanctonum sanctonum*) despite the presence of cats, mongooses, rats, and humans.

*Alsophis rijgersmai* occurs on Anguilla and St. Barthelemy, where Sadjak and Herderson (1990) conducted two surveys of six and three days and observed three and five snakes respectively. The species is also presumed to occur on Scrub Island off the eastern coast of Anguilla. St. Martin/St. Maarten has a dense mongoose population and racers are thought to have been extirpated; the most recent report of *A. rijgersmai* on this island is from 1951 (Brongersma, 1959).

Daltry *et al*. (1997) reported ‘high densities’ of *A. rufiventris* on Saba and St. Eustatius, especially in upland areas, but warned of the risk of complacency due to the islands’ small sizes and fragile ecosystems. During field research on St. Eustatius in March 1997, Daltry *et al*. captured and measured 40 (35 male; five female) live racers. Based on interactions with local residents, the authors suggested that the species was already declining on St. Eustatius as a result of habitat loss, human persecution, and predation by black rats and domestic cats. The racer’s population density was not estimated for either island, but was said to be ‘very abundant’ within its current distribution range, with robust populations on both islands. The species appeared to be more abundant on Saba than St. Eustatius (e.g., one racer was detected every thirty minutes in the Saban rainforest, whereas sightings on St. Eustatius occurred ‘much less frequently’; Daltry et al., 1997). On St. Eustatius, racers appeared to be most abundant in the Quill National Park (Daltry et al., 1997). From 3 - 23 June 2004, Savit *et al*. (2005) encountered 182 racers along a single hiking trail on the western slope of the Quill. The authors determined that the highest racer encounter rates were along sections of the trail that contained the most rocks. Rocks may be attractive to snakes due to the availability of refuges or because prey species may refuge in cavities surrounding rocks, *A. schwartzi* (Lazell, 1972), was closely associated with rocks in the Quill. From September to December 2016, Zobel (2016) surveyed three hiking trails in the Quill where she captured, recaptured and PIT-tagged 68 racers (36 male; 32 female).

*Alsophis sibonius* (Cope, 1879), formerly considered a subspecies of *Alsophis antillensis,* is endemic to Dominica. During a seven-day survey of the island, Sadjak and Henderson (1991) found nine racers in Cabrits peninsula. In 2008, White *et al*. (2008) encountered 19 snakes based on 267 min of focal animal observations in a survey of snake activity and habitat associations. Sadjak and Henderson (1991) observed two *A. a. manselli* racers in a four-day survey on Montserrat in 1987. Local residents were said to be familiar with the snake, and mongooses were not observed or reported.

*Borikenophis* (formerly *Alsophis*) *portoricensis* (Reinhardt and Lütken, 1862) inhabits the Puerto Rican Bank, including Puerto Rico mainland and the Virgin Islands. MacLean (1982) reported the species to be rare or extirpated on the larger Virgin Islands but still moderately common in Puerto Rico and many smaller islands (Barun et al., 2007). Total snake encounters over a five-year study period (2001-2005) numbered 205, however population abundance or density was not estimated (Barun et al., 2007).

Introduced and invasive mammalian predators such as rats, cats, dogs and mongooses pose a threat to most species of racers (Seaman and Randall, 1962; Henderson, 1992). This is the case for *A. rufiventris* on St. Kitts and Nevis, which is thought to have been extirpated by the mongoose following its introduction circa the early 1900s (Sajdak and Henderson, 1991; Borroto-Páez and Woods, 2012). Consequently, the current range of *A. rufiventris* in the Lesser Antilles is now limited to the two smallest islands of the Caribbean Netherlands: Saba and St. Eustatius. These last remaining habitats represent just 10.9% of the racer’s original range of 302 km^2^ (Fig. 1; Sajdak and Henderson, 1991). Besides invasive species, other external factors that contribute to population declines include volcanic activity (Young and Ogrodowczyk, 2008), hurricanes (Daltry, 1991; 2006; Zobel et al., 2018), development (Daltry et al., 2017), road kills by traffic (Hypolite et al., 2007), and habitat loss and alteration (Henderson and Tolson, 2006).

**Figure 1.**
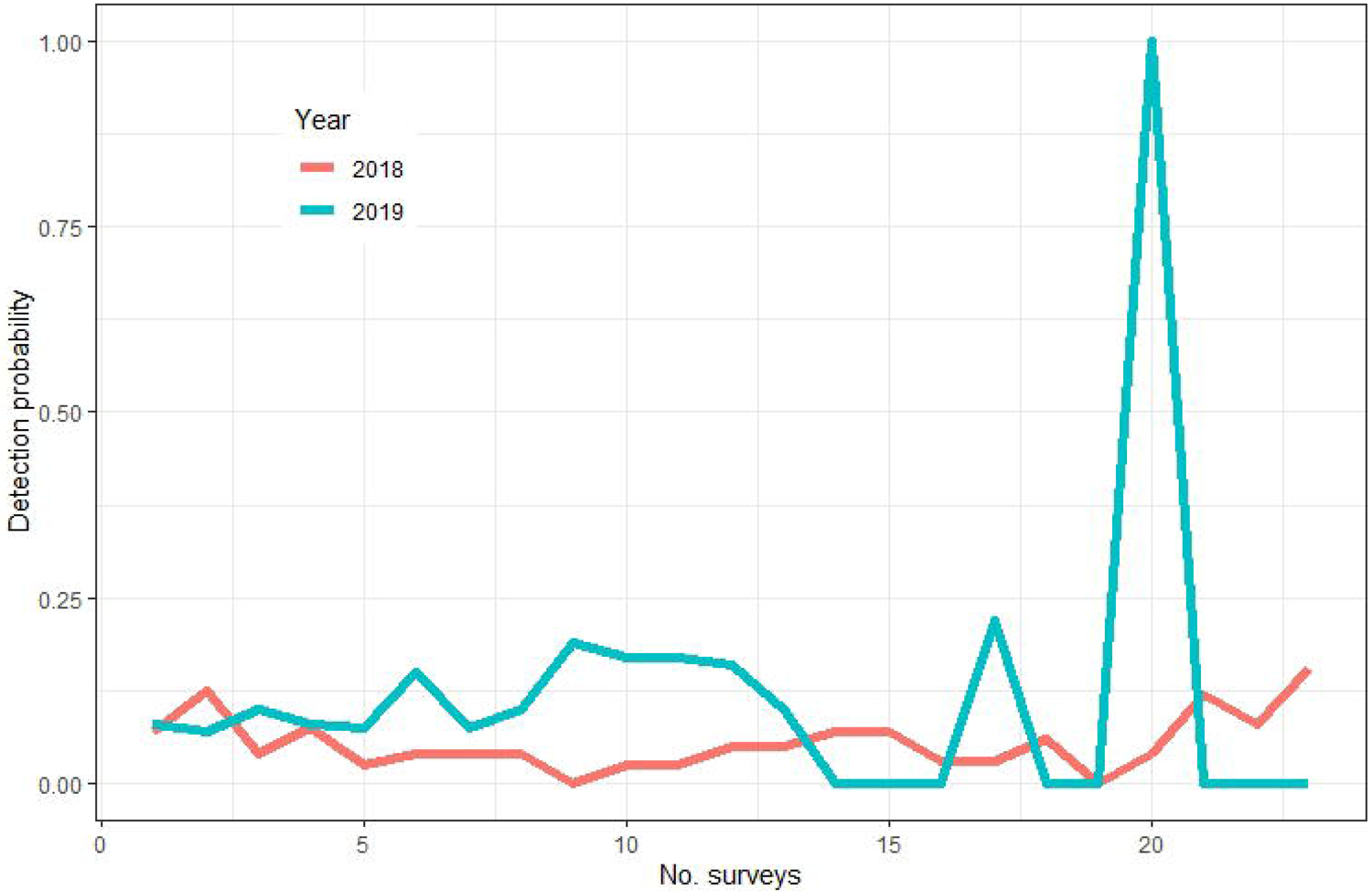
Distribution of *Alsophis rufiventris* on the St. Christopher and Saba banks. Dots indicate locality records, X’s mark extirpated populations on St. Kitts and Nevis, and red stars mark fossil localities (adapted from Maley et al., 2006).

### Hurricane impacts

Rising sea surface temperatures are predicted to cause more frequent and intense hurricanes in the Caribbean (Webster et al., 2005; Biasutti et al., 2012). The 2017 Atlantic hurricane season recorded six major hurricanes, including hurricanes Irma (cat. 5, maximum wind speed 285 km h−1) and Maria (cat. 5, max. wind speed 280 km h−1; www.nhc.noaa.gov/). In addition to the natural and anthropogenic disturbances mentioned above, major hurricanes can increase the risk of extinction of Caribbean reptile species - especially those with restricted distributions and small, closed populations (Powell and Henderson, 2005). Our research was conducted prior to and following two major hurricanes which impacted St. Eustatius in 2017, causing extensive damage to forest vegetation (Eppinga and Pucko, 2018). Insular populations that have evolved in the absence of mainland predators and competitors are considered especially vulnerable (Powell, 2006). Whereas the critically endangered Lesser Antillean iguana (*Iguana delicatissima* Laurenti, 1768) has been the focus of extensive conservation efforts on St. Eustatius over the past few years (Debrot et al., 2013; van den Burg et al., 2018; van Wagensveld and van den Burg, 2018), the red-bellied racer has received less attention (Powell, 2006). This was in part due to the belief that the species was thriving, especially in the Quill (HM, pers. obs.). On a hike to the crater rim (∼400 a.s.l), casual hikers would regularly encounter snakes and the population was thought to be stable. However, many island species are in various stages of decline, and less enigmatic reptile species have been ignored by even professional conservation biologists (Powell, 2006). Despite its current classification under IUCN criteria as Vulnerable (Daltry and Powell, 2016), (in 1996 it was classified as Endangered; Daltry et al., 1997), the population was potentially impacted following two major hurricanes in September 2017. In many cases, as evidenced earlier in this paper, baseline data are lacking. Whilst some preliminary studies have been conducted on *A. rufiventris* on St. Eustatius (e.g., Daltry et al., 1997; Savit et al., 2005; Zobel et al., 2018), we present the first quantitative assessment of the population. The results of this study can be used as a baseline to monitor long-term population trends and survival of the racer in the context of environmental and ecological changes.

Occupancy is defined as the probability of a site being occupied by a species (MacKenzie et al., 2002); fluctuations in occupancy estimates provide a robust proxy for population stability. Such estimates are particularly useful for sparsely populated or cryptic species that are difficult to detect (Beaudrot et al., 2016). The goal of our study was to estimate abundance, detection probability and site occupancy of the red-bellied racer at three different time periods. We hypothesized that if the racer’s population size was stable based on surveys conducted prior to and post hurricanes, this would suggest that snakes are resilient to changes. If the racer’s abundance estimates varied, however, this would suggest that external factors (direct and indirect) are impacting life history characteristics and vulnerability of the species. An additional goal of our study was to determine the influence of specific geographic and ecological covariates on site occupancy, abundance and detection probability and abiotic/biotic variables which may be correlated with the presence of the racer.

## METHODS

### Study Area

Our study took place on St. Eustatius (17°28’N, 62°57’ W), a small (21 km^2^) island with a human population of approximately 3,900 (Statistics Netherlands, 2018) located approximately 12.5 km north-west of St. Kitts (Fig. 1). Temperature typically ranges from 25 to 33°C. Average annual rainfall is 986 mm, with upper elevations of the Quill receiving 1,500 – 2,000 mm per year (Rojer, 1997; van Andel et al., 2016). Our study took place within and outside the boundaries of the Quill (∼220 ha) and Boven National Parks (∼320 ha). Vegetation in the Quill ranges from evergreen seasonal secondary forest inside the crater and higher slopes to thorny woodland below 250 meters (Stoffers, 1956). Vegetation in Boven comprises dry forest fragments with areas of open, grassy shrubland (van Andel et al., 2016).

### Species Description and Behavior

The red-bellied racer (hereafter ‘racer’) is a moderately sized (max total length: male = 1179mm; female = 976 mm (Daltry et al., 1997); male = 1046 mm; female = 1070 mm (this study)), sexually dimorphic colubrid (Maley et al., 2006). Racers are variable in color and pattern, and members of the same species may vary morphologically on different islands. A diurnal species, the racer forages through leaf litter in search of lizards (*Anolis* spp. Daudin, 1802), frogs (*Eleutherodactylus johnstonei* Barbour, 1914), iguana (*Iguana delicatissima* Laurenti, 1768) hatchlings and the occasional egg (Powell et al., 2005). Despite growing to a length of approximately one meter, it is considered a small snake that is harmless to humans. If captured, however, it will release an unpleasant-smelling cloacal secretion (Malhotra and Thorpe, 1999). Nevertheless, the racer is often persecuted by people who fear them (Powell, 2006) and encounters with humans can result in the demise of the snake. While the racer’s venom is capable of subduing its prey, its small size and rear-facing fangs prevent it from attacking larger victims. Little is known about *Alsophis’* breeding habits, however a captive breeding program at Jersey Zoo resulted in *Alsophis antiguae* producing 11 eggs, of which five hatched (Daltry et al., 2001).

The racer shows remarkable sexual dimorphism, whereby males and females can be reliably distinguished (Daltry et al., 1997). Males are predominantly chocolate brown to black on the dorsum with conspicuous paler brown/yellow blotches; females are predominantly grey to medium brown dorsally with darker streaks (Daltry et al., 1997). Additionally, female racers typically have longer and broader heads than males. Male racers can also be distinguished by the higher number of pairs of subcaudal scales. The tails of adult males are also noticeably thickened at the base and relatively long, accounting for 26.4–30.9 % of the snake’s total body length (Daltry et al., 1997).

Racers are found in a variety of natural and altered habitats, ranging from thorny woodland to moderately mesic hillsides. The species was previously described as ‘thriving’ by Maley et al. (2006), being common even in close proximity to human activity and habitation. The Quill and Boven National Parks are considered important habitats for the racer (Daltry et al., 1997). Besides the introduced brahminy blind snake (*Indotyphlops braminus* Daudin, 1803; Snyder et al. 2019), the only other native snake species that occurs on St. Eustatius is the Leeward Blindsnake (*Antillotyphlops geotomus* Thomas, 1966; van Wagensveld et al. 2020). *A. rufiventris* has few natural predators but has been observed being consumed by American Kestrel (*Falco sparverius* Linnaeus, 1758; HM pers. obs.). However, the species is threatened by non-native species such as free-roaming cats, dogs, chickens (*Gallus gallus domesticus* Linnaeus, 1758), and black rats (Henderson, 1992). Black rats are thought to attack racers, especially when rats are hungry or defending young (Daltry et al., 1997). The racer is too small to be able to consume adult rats (Daltry et al., 1997). Attacks on the snake, on the other hand, may have implications for the reproductive success of racers, which can lose portions of their tail (where the hemipenis and retractor muscles are located; Daltry, 2006). Daltry et al. (1997) reported 55% of captured snakes with incomplete tails, indicative of likely heavy predation pressure on St. Eustatius and Saba. House mice (*Mus musculus* Linnaeus, 1758), while present on St. Eustatius (Madden et al., 2019), are not thought to constitute a threat to the snake. The presence of free-roaming herbivores in the Quill and Boven (Madden, 2020) could indirectly impact racers through trampling or loss of vegetation through grazing (McCauley et al., 2006), however this has not been studied. Fortunately the island is currently not occupied by the mongoose, whose introduction to other Caribbean islands is thought to be responsible for numerous reptile extirpations (Table 1; Henderson, 1992; Powell and Henderson, 2005; Powell et al., 2005).

**Table 1.** Overview of all islands in the Lesser Antilles where *Alsophis* sp. is known to have been present, or is currently present, and the presence or absence of mongoose. On at least five islands *Alsophis* has been extirpated and in all cases the presence of mongoose is known (Powell and Henderson, 2005). Adapted from Powell and Henderson (2005).

| Island | Mongoose | *Alsophis* |
| --- | --- | --- |
| Anguilla | No | Yes |
| Antigua | Yes | No |
| Dominica | No | Yes |
| Great Bird | No | Yes |
| Guadeloupe | Yes | No |
| Marie-Galante | Yes | No |
| Montserrat | No | Yes |
| Saba | No | Yes |
| Scrub | No | Yes |
| St. Barthelemy | No | Yes |
| St. Christopher & Nevis | Yes | No |
| St. Eustatius | No | Yes |
| St. Martin/St. Maarten | Yes | No |
| Terre-de-Bas | No | Yes |
| Terre-de-Haut | No | Yes |

### Surveys: Pre-hurricane Surveys

Specific survey locations on the Quill include four areas: NW facing slope (range 238-350 m.a.s.l.), SW facing slope (range 220-400 m.a.s.l.), and East facing slope/Botanical Garden (range 106-220 m.a.s.l.) and inside the crater (range 300-351 m.a.s.l.). Location of each area surveyed is indicated in Figure 2. At each site, snakes were located using transects along the trail (10 m wide on each side) and/or perpendicular to the trail in the NW and SW areas and inside the crater; for these three locations, eight people survey in parallel transects, at 6 m apart (50 m wide total) and 350 long, covering an area of 1.7 ha each (see Online Supporting Information Table S1 for specific areas). Surveys commenced after 9:00 h and finished around 14:00 h, between 21st January 2011 and 24th January 2011. When a snake was detected, a parallel point was selected at an equivalent position from the center of the transect, these are called null points thereafter. At each point (snake and null), all measurements described below were conducted. We calculated snake density by dividing the number of individuals detected by the calculated area of each transect.

**Figure 2.**
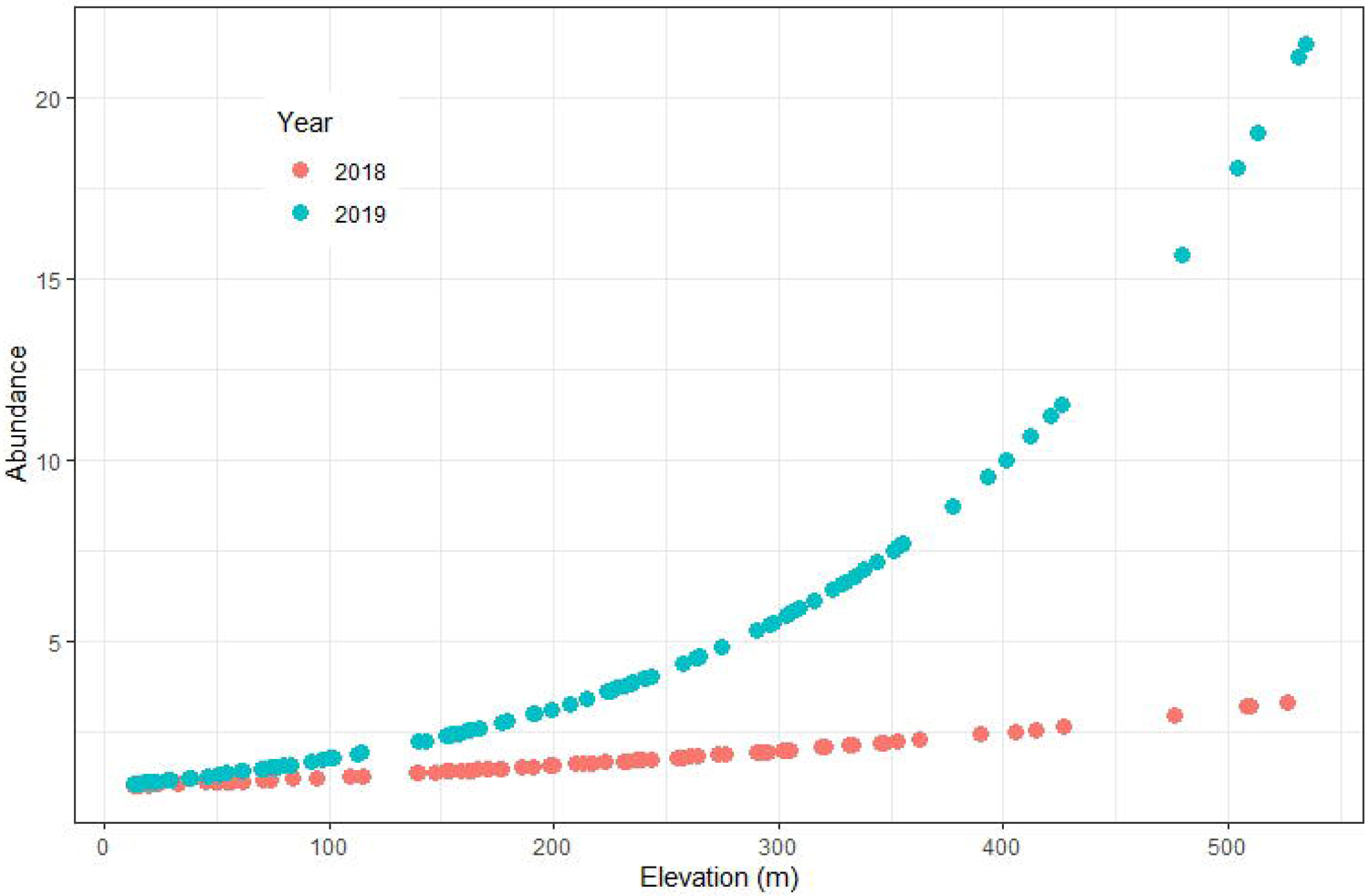
Map of St. Eustatius showing 2011 survey areas (yellow stars), 2018 and 2019 transects (red lines), 20m contours (black lines) and vegetation types. Adapted from de Freitas et al. (2014).

### Environmental Measurements (Pre-hurricane)

On each point were a snake was detected or the null point, we conducted the following measurements: air temperature and relative humidity with a hand-held meter; hemispherical photography pointing above the place, with a digital camera with a fish-eye lens, to use it to calculate leaf area index and canopy cover (HemiView 2.1 software, Delta-T/Dynamax); photography of the area enclosed in a 1m^2^ frame placed on the forest floor, to use it to measure the fractions of bare soil, green vegetation, leaflitter, rocks and wood; depth of the litterfall using a ruler and measuring five points inside the frame; location and elevation with a GPS receiver.

### Post-hurricane Surveys

In 2018 and 2019 we surveyed racers in the Quill and Boven National Parks (total survey area 11.42 ha). Surveys were conducted from December 2017 to May 2018, and from March to June 2019. To maximize detection and minimize disturbance, all surveys were conducted along existing hiking trails. Using the existing network of trails allowed us to survey snakes in an otherwise steep topographical environment with, at times, rocks or dense vegetation. We conducted surveys along a 100-meter line transect by looking and listening for *A. rufiventris* while walking at a very slow pace. All surveys were conducted between 0700 and 1900 hrs. Once detected, the perpendicular distance of the snake from the center of the transect was measured with a tape measure.

### Morphological Measurements and PIT-tagging of Snakes

We measured morphological characteristics of 60 racers (20 in 2018 and 40 in 2019; snout – vent; vent – tail; total length, cm) and weighed individuals using a pesola scale (g). We scanned all captured racers for a PIT tag during surveys in 2018 and 2019. In 2019, if no PIT tag was detected in the captured snake we inserted one (10.9 x 1.6 mm) using a 2mm sterile needle (Online Supporting Information Table S2). After sterilizing the skin, the PIT tag was placed subcutaneously in the mid-body region on the mediolateral side. Markings on the body and other morphological differences described earlier enabled us to confidently determine the sex of captured individuals.

**Table 2.** Top models for the presence of snakes, AICc, corrected Akaike Information Criterion, delta = change in AICc as compared to the best model, weight = the weight of the model as compared to all other models in the analysis. The last line in italics represents the worst model.

| **%**<br>**bare**<br>**soil** | **%**<br>**green**<br>**vegetat**<br>**ion** | **%**<br>**litter** | **%**<br>**wood** | **Leaf**<br>**Area**<br>**Index** | **Litter**<br>**depth** | **Soil**<br>**temp** | **df** | **logLik** | **AICc** | **ΔAICc** | **weight** |
| --- | --- | --- | --- | --- | --- | --- | --- | --- | --- | --- | --- |
| -3.933 | NA | NA | NA | NA | -0.022 | 0.057 | 3 | -90.75 | 187.69 | 0 | 0.07 |
| -3.343 | NA | NA | NA | 0.238 | -0.017 | NA | 3 | -90.8 | 187.77 | 0.083 | 0.067 |
| -3.957 | NA | NA | NA | 0.143 | -0.024 | 0.035 | 4 | -89.95 | 188.21 | 0.516 | 0.054 |
| -3.188 | NA | NA | 1.269 | 0.206 | -0.017 | NA | 4 | -90.09 | 188.49 | 0.796 | 0.047 |
| -4.442 | NA | -1.06 | NA | NA | -0.02 | 0.086 | 4 | -90.25 | 188.80 | 1.108 | 0.04 |
| -4.541 | NA | -1.23 | NA | 0.159 | -0.021 | 0.066 | 5 | -89.28 | 189.02 | 1.33 | 0.036 |
| -3.273 | 1.223 | NA | NA | 0.229 | -0.018 | NA | 4 | -90.57 | 189.45 | 1.756 | 0.029 |
| -3.711 | NA | NA | 0.679 | NA | -0.021 | 0.05 | 4 | -90.60 | 189.51 | 1.817 | 0.028 |
| -3.846 | 0.854 | NA | NA | NA | -0.023 | 0.054 | 4 | -90.65 | 189.60 | 1.908 | 0.027 |
| NA | 0.657 | -0.6 | NA | 0.12 | NA | -0.004 | 4 | -95.39 | 199.08 | 11.39 | 0 |

### Ecological and Geographic Covariates

The following covariate data were collected: elevation (m): measured with a handheld GPS; temperature (°C): measured with a temperature gauge; cloud cover: estimated by eye, categorical (0–4; 0= no cloud, 4 = full cloud); habitat: 6 habitats were selected based on vegetation descriptions by de Freitas et al., (2014; Fig. 1); rainfall: total monthly rainfall from the previous month was included in the dataset following the completion of surveys (data downloaded from http://www.seawf.com/rainhist2.php).

### Habitat descriptions

We split our survey areas into six habitat types based on vegetation descriptions by de Freitas et al. (2014; Fig. 2): 1. *Pisonia-Justicia* and *Pisonia-Bothriochloa* hills (H1, H2). Elevation: 13 - 177m; 2. *Pisonia-Eugenia* mountains (M5). Elevation: 155-354 m; 3. *Pisonia-Eugenia, Chionanthus-Nectandra* and *Capparis-Antirhea* mountains (M5, M3b, M6).

Elevation: 275-354 m; 4. *Myrcia-Quararibea* (M1). Elevation: 296-400 m; 5. *Coccoloba-Chionanthus* and *Chionanthus-Nectandra* mountains (M2, M3a). Elevation: 393-534 m; 6. *Capparis-Pisonia* mountains (M4). Elevation: 80-275 m.

## STATISTICAL ANALYSES

### Pre-hurricane Methods

#### Abiotic and Biotic Variables Influencing Detection

To assess potential relationships between the presence and absence of *Alsophis* (where a snake was located or not) correlated with biotic and abiotic variables we used a model selection approach with a generalized linear model (GLM). Variables evaluated included leaf litter depth, percentage of bare soil, percentage of litter, percentage of green vegetation, percentage of wood, soil temperature, above-ground air temperature (30 cm), and leaf area index above the site where the snake or control site was located. Data for the control site (null point) were collected at a position in the other half of the transect equivalent to the one where the snake was found. These data were only collected for 2011 surveys. The biotic and abiotic data for 71 sites where snakes were located and 75 control sites were collected.

We tested for collinearity using kendall’s correlation analysis. We performed pairwise correlations, whereby all correlations were below ± 0.39 except air and soil temperature (cor = 0.49), thus we eliminated air temperature from further analysis. We performed a GLM analysis with a binomial link (snake presence or absence; Crawley, 2007; Weigelt & Kreft, 2013). Models were ranked according to the corrected Akaike’s Information Criterion (AICc) and the model with the lowest AICc is considered the best-fit model (Akaike, 1974; Sugiura, 1978). Those models with AICc differences <2 relative to the best-fit model were also considered as plausible (Arnold, 2010). The 95% normal confidence intervals of the coefficients are calculated from the most parsimonious models. All GLM analyses were performed in *R* (R Development Core Team, 2013) using the *MASS* package (Venables & Ripley, 2002) and the *dredge* function in the *MuMIm* package (Barton, 2019) for comparing all models.

#### Post-hurricane Methods

Density estimation: We used the Schnabel index to estimate population size based on marked and recaptured snakes in 2019, and compared this with marked/recaptured snakes from 2016 (Zobel, 2016). We used a likelihood-based, single-season occupancy model (MacKenzie et al., 2017) in the package ‘wiqid’ (Meredith, 2020) to estimate site occupancy (ψ) in relation to habitat and elevation, while accounting for detectability (*p*). During wildlife surveys, individuals can go undetected, either because they are truly absent at the sampling site, or because they are present, but were not observed. Visiting each sampling site on multiple occasions enables the estimation of detection probability, which can be incorporated into occupancy modeling to correct for imperfect detection (MacKenzie et al., 2002). Occupancy models link a state model determining occupancy at each site with an observation model for detection, which is conditional on occupancy. Occupancy models were based on three assumptions: (i) the racer population in each site was closed to birth, death, immigration, and emigration during the sampling period; (ii) counts at each site were independent; and (iii) individuals were not double counted within a single sampling occasion (Murray and Sandercock, 2020). We first modeled detection probability and then occupancy probability. We subsequently used a built-in covariate function which estimated detection in relation to whether the species had previously been detected or not. The function ‘occSStime’ allowed us to include a time trend in our analysis based on survey effort. The function ‘occSSrn’ (Royle and Nichols, 2003) provided an estimation of site occupancy (ψ), average number of individuals per sample, lambda (λ) and sampling detection probability (*p*). Finally, we repeated some of the analyses with the package ‘unmarked’ (Fiske and Chandler, 2011) using a basic N-mixture model to estimate racer abundance (λ) and detectability (*p*). This allowed us to test the influence of covariates such as rainfall, temperature, week of the year on abundance estimates. Quadratic terms for all variables were also included to account for the possibility of non-linear responses. We ranked the resulting models using Akaike’s Information Criterion, corrected for small sample size (AICc; Burnham and Anderson, 2002) in the package ‘MuMIn’ (Barton, 2020) where the detection model with the lowest AICc is best-fitting. All analyses were performed in the R environment version 4.0.3.

## RESULTS

### Presence of *Alsophis* with Biotic and Abiotic Variables

A total of 128 different models were evaluated and eight models had ΔAICc ≤2 with a total weight of 0.398 (Table 2). The only variables that were consistent among all eight models was litter depth and fraction of bare soil, which were negatively correlated with the presence of snakes. Leaf Area Index (LAI) and the soil temperature were both positively correlated with the presence of snakes. All seven variables were included in at least one of the models. The average coefficients of the best models resulted in wide confidence intervals in four of the seven variables which were included in the models. Evaluating the confidence intervals as estimated from the multiple models only two variables were consistent in that they excluded zero (the null model), fraction of bare soil and litter depth, both of which had a negative effect on snake occurrence (Fig. 3).

**Figure 3.**
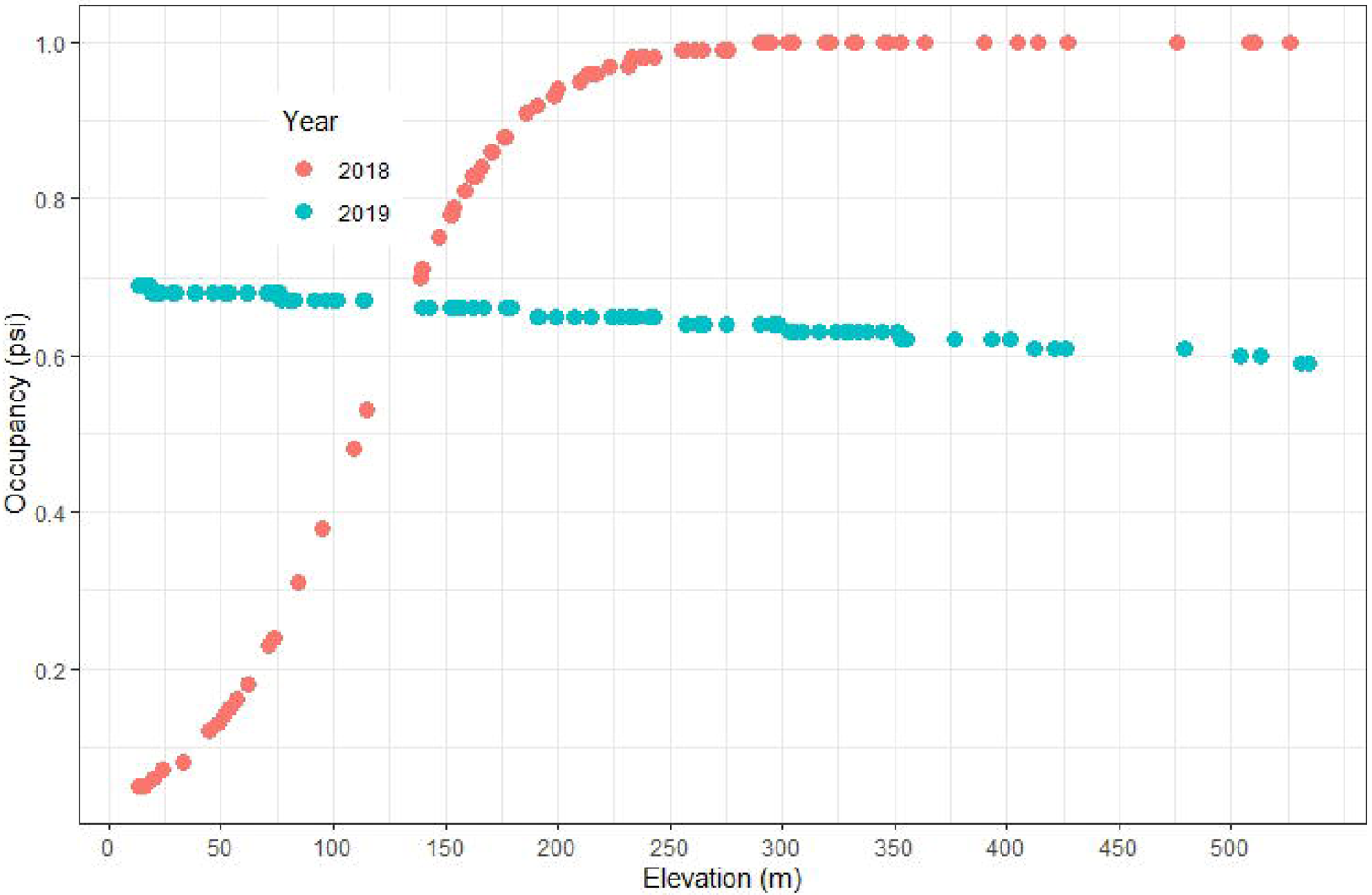
Mean coefficients of the selected models and their 95% confidence intervals. Confidence intervals are estimated from variation of the coefficient among the eight most parsimonious models (selected models). Vertical line represents a coefficient of zero.

### Pre- and post-hurricane Survey Densities and Encounter Rates

Pre-hurricane snake densities/ha were higher than post-hurricane densities across all comparable habitats. For example, in the Quill crater, pre-hurricane densities varied between 11.9 (min 7.5 - max 18.7) and 22.4 (min 9.1 - max 51.3) snakes/ha, whereas post-hurricane densities varied between 0 (min 0 - max 4.9) and 2.1 (min 0.4 - max 8.4) snakes/ha (Fig. 4). Similarly, in the Botanical Garden/Around the Mountain S, densities decreased from 13.2 (min 7.3 - max 23.1) snakes/ha in 2011, to 3.3 (min 1.1 - max 9.2) snakes/ha in 2018 and 0 (min 0 - max 3.9) snakes/ha in 2019 (Fig. 4). We excluded the northern hills, where we only found one snake, and the Quill rim, which we did not survey in 2011 (Figure 3). In 2011 (pre-hurricane), based on 146 transect surveys (21-24 January), we encountered 64 racers. Post-hurricane, we detected 54 individuals in 2018 (based on 1,468 transects) and 60 in 2019 (based on 1,084 transects). This translates to a raw encounter rate across all habitats surveyed of 16.0 snakes/hour in 2011, 0.34 snakes/hour in 2018 and 0.41 snakes/hour in 2019.

**Figure 4.**
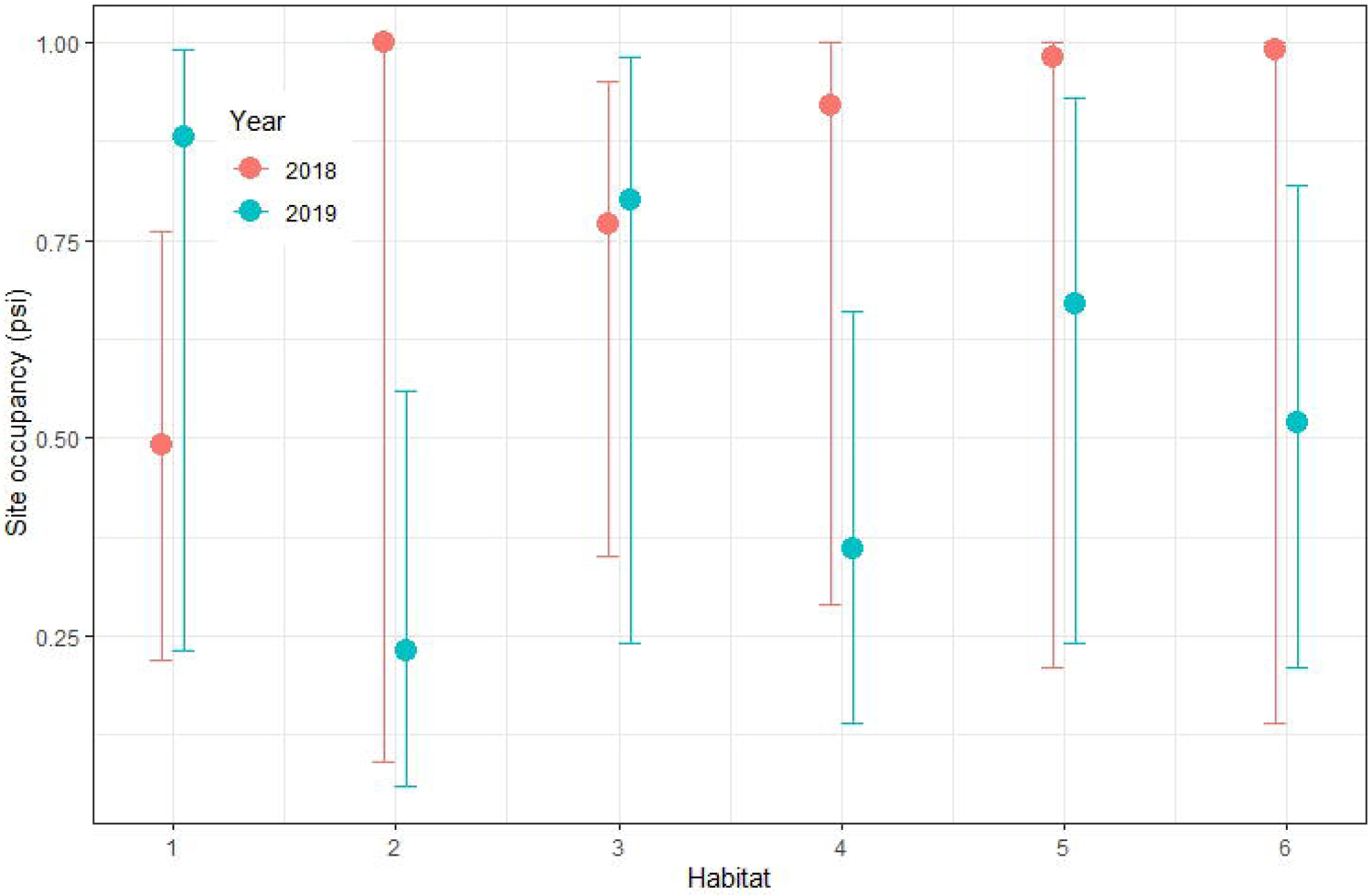
Estimated density (± SE) per hectare of *Alsophis rufiventris* on St. Eustatius based on surveys conducted in 2011 (left), 2018 (middle) and 2019 (right). Note that in 2011 the Northern Hills (H1-H2) and Quill rim (M5) were not surveyed.

### Post-hurricane Survey Population Size

We conducted surveys in 2018, following Hurricanes Irma and Maria, and in 2019, surveying an area of 6.6 ha in the Quill National Park and 4.8 ha in Boven National Park. Average transect length was 100.73 ± 0.35 m. Line transects were repeated a minimum of two and a maximum of 24 times at different sites. Average transect time was 7.08 ± 0.11 minutes. Estimates of population size varied among sites surveyed. Using data from PIT-tagged snakes and based on the Schnabel index estimate of marked and recaptured snakes, we calculated a population size of 44.39 (± 0.46) in 2016 based on a survey area of 3.5 ha in the Quill. For 2019 we calculated a population size of 310.5 (± 0.16) based on a survey area of 6.6 ha in the Quill, and 249 (± 0.18) based on a survey area of 4.8 ha in Boven (Table 3).

**Table 3.** Overview of 2018 and 2019 site occupancy (ψ; the probability of a site to be occupied) and mean abundance (*λ;*; average individuals per sample) estimates per habitat (with upper and lower confidence intervals and SD) of *Alsophis rufiventris* on St. Eustatius. 1 = Northern hills; 2 = Quill main trail; 3 = Quill Around the Mountain S (upper); 4 = Quill crater; 5 = Quill rim; 6 = Botanical Garden/Around the Mountain S (lower). Black text = wiqid; red text = unmarked.

| Year | Habitat | $\psi$ | low CI | upp CI | $\lambda$ | SD |
| --- | --- | --- | --- | --- | --- | --- |
| 2018 | 1 | 0.488<br>0.806 | 0.224<br>0.374 | 0.759<br>0.997 | 0.264 | 0.168 |
| 2018 | 2 | 0.774<br>0.966 | 0.355<br>0.568 | 0.955<br>1.000 | 1.785 | 0.288 |
| 2018 | 3 | 0.925<br>0.850 | 0.289<br>0.422 | 0.997<br>0.999 | 2.001 | 0.189 |
| 2018 | 4 | 0.978<br>0.888 | 0.208<br>0.467 | 1.000<br>1.000 | 2.011 | 0.102 |
| 2018 | 5 | 0.994<br>0.920 | 0.141<br>0.507 | 1.000<br>1.000 | 2.865 | 0.355 |
| 2018 | 6 | 0.998<br>0.946 | 0.092<br>0.540 | 1.000<br>1.000 | 1.544 | 0.227 |
| 2019 | 1 | 0.883<br>0.973 | 0.229<br>0.237 | 0.995<br>1.000 | 1.631 | 0.515 |
| 2019 | 2 | 0.798<br>0.655 | 0.237<br>0.074 | 0.981<br>1.000 | 5.062 | 1.920 |
| 2019 | 3 | 0.675<br>0.941 | 0.235<br>0.195 | 0.933<br>1.000 | 6.548 | 0.922 |
| 2019 | 4 | 0.521<br>0.743 | 0.211<br>0.097 | 0.816<br>1.000 | 6.181 | 0.876 |
| 2019 | 5 | 0.364<br>0.891 | 0.145<br>0.158 | 0.659<br>1.000 | 14.045 | 5.403 |
| 2019 | 6 | 0.231<br>0.824 | 0.065<br>0.125 | 0.565<br>1.000 | 3.246 | 1.114 |

### Abundance estimates

Estimated racer densities per hectare in 2011, 2018 and 2019 mainly varied in their differences among habitat types but also varied greatly in the dispersion of the estimates among years (Fig. 4). The naïve occupancy (i.e., proportion of sites the species was detected; ψ) of racers was 84% (48% - 99% CI) in 2018, and 80% (32% - 99% CI) in 2019. Racer occupancy increased with increasing elevation in 2018, but decreased with increasing elevation in 2019. Occupancy varied across the six habitats sampled, but exhibited similar patterns in 2018 and 2019 (Figs. 5a and 5b). Site occupancy and mean abundance estimates per habitat for 2018 and 2019 are provided in Table 5. Mean detection probability (*p*) per transect was 0.05 (0.03 - 0.07 CI) in 2018, and 0.07 (0.05 - 0.11 CI) in 2019 (package: ‘wiqid’). In 2018, *p* was 0.02 (0.01 - 0.06 CI), and 0.03 (0.01 - 0.12 CI) in 2019 (package: ‘unmarked’). During 2018 surveys, *p* increased from 0.05 (0.03 - 0.07 CI) to 0.09 (0.03 - 0.21 CI) with a previous racer detection. During 2019 surveys, *p* did not increase with a previous detection. The number of surveys was not an important predictor of *p* in 2018, however *p* increased significantly in 2019 with more (15 - 20) visits (Fig. 6). Estimated mean abundance (λ) per transect was 1.82 (0.66 - 5.01 CI) in 2018, and 1.60 (0.39 - 6.65 CI) in 2019 (package: ‘wiqid’). Estimated λ was 1.90 in 2018 and 4.50 in 2019 (package: ‘unmarked’). When habitat and elevation were included as fixed effects, mean λ was 1.72 (SD ± 0.52) in 2018 and 4.69 (SD ± 4.16) in 2019. This translates to a population estimate across the entire study area (11.42 ha) of 139.40 in 2018 and 506.16 in 2019. Snake abundance increased with elevation, however the observed rate of increase among years was different, showing a drastic increase in 2019, while the rate of increase in 2018 was small (Fig. 7). Occupancy rates varied dramatically between the two years, showing an occupancy rate of 80% or more at elevations of 150m in 2018, while the occupancy rate at all elevations was more or less equal at all elevations (±65%; Fig. 8). Site occupancy in 2018 ranged from 0.49 (0.22 - 0.76 CI) in the northern hills to 0.99 (0.14 - 1.00 CI) along the crater rim of the Quill, though we note the very large confidence intervals. Conversely, occupancy in 2019 ranged from 0.23 (0.07 - 0.57 CI) in the botanical garden/lower Quill slopes to 0.88 (0.23 - 1.00 CI) in the northern hills.

**Figure 5a.**
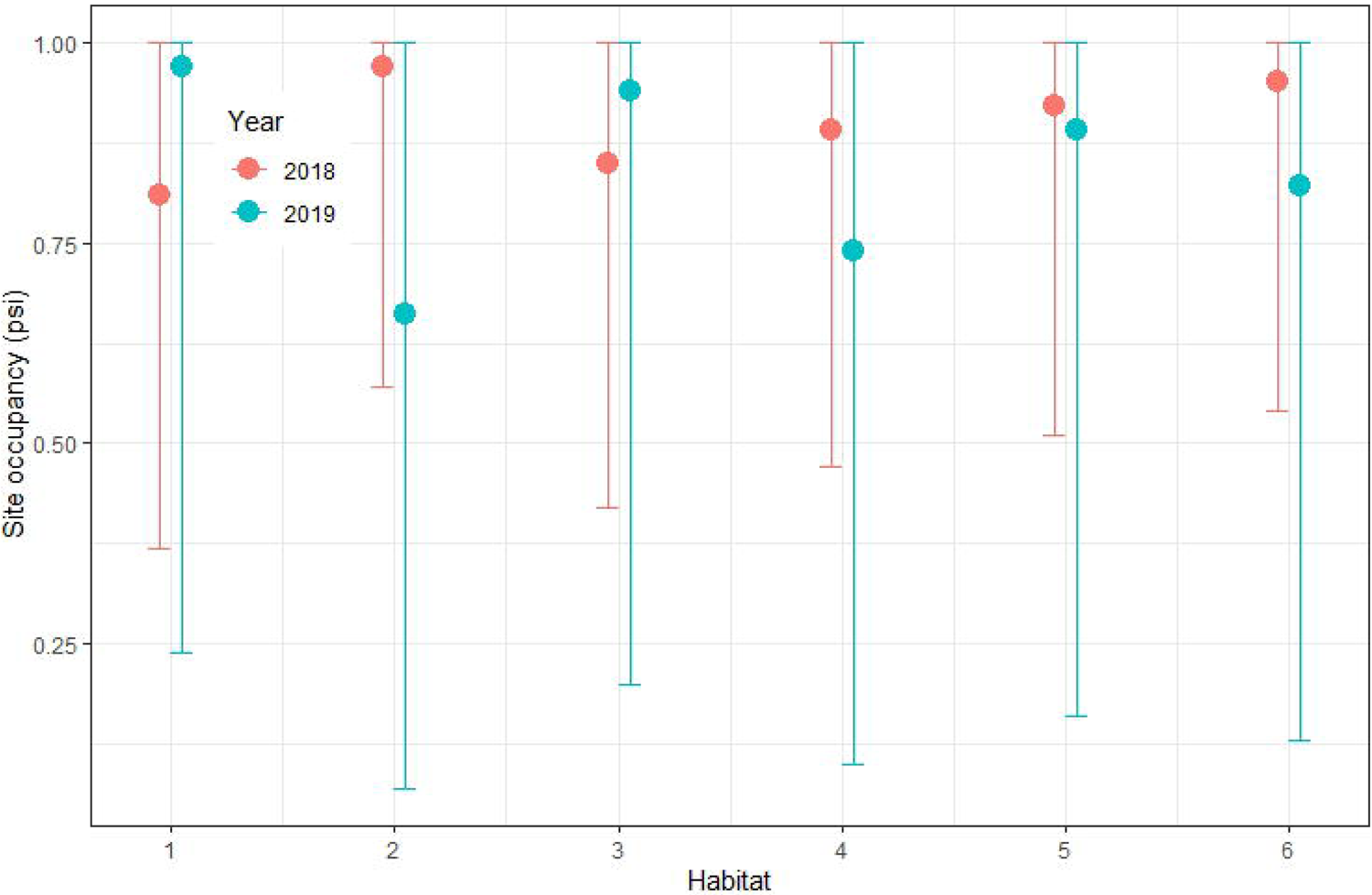
Mean abundance (± SE) per transect per habitat of *Alsophis rufiventris* on St. Eustatius based on surveys conducted in 2018 and 2019. 1 = Northern hills; 2 = Quill main trail; 3 = Quill Around the Mountain S (upper); 4 = Quill crater; 5 = Quill rim; 6 = Botanical Garden/Around the Mountain S (lower). Package: wiqid.

**Figure 5b.**
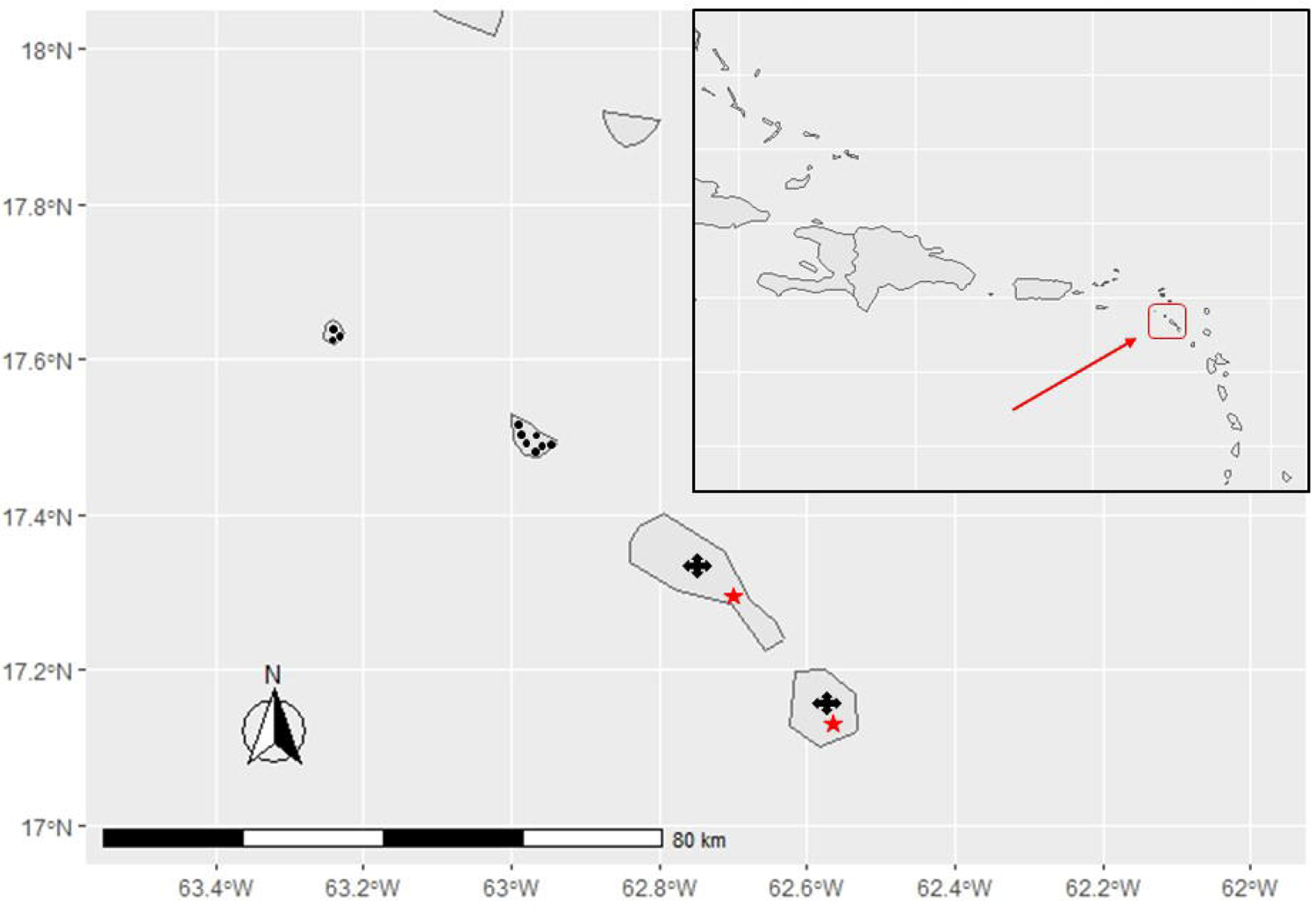
Mean abundance (± SE) per transect per habitat of *Alsophis rufiventris* on St. Eustatius based on surveys conducted in 2018 and 2019. 1 = Northern hills; 2 = Quill main trail; 3 = Quill Around the Mountain S (upper); 4 = Quill crater; 5 = Quill rim; 6 = Botanical Garden/Around the Mountain S (lower). Package: unmarked.

**Figure 6.**
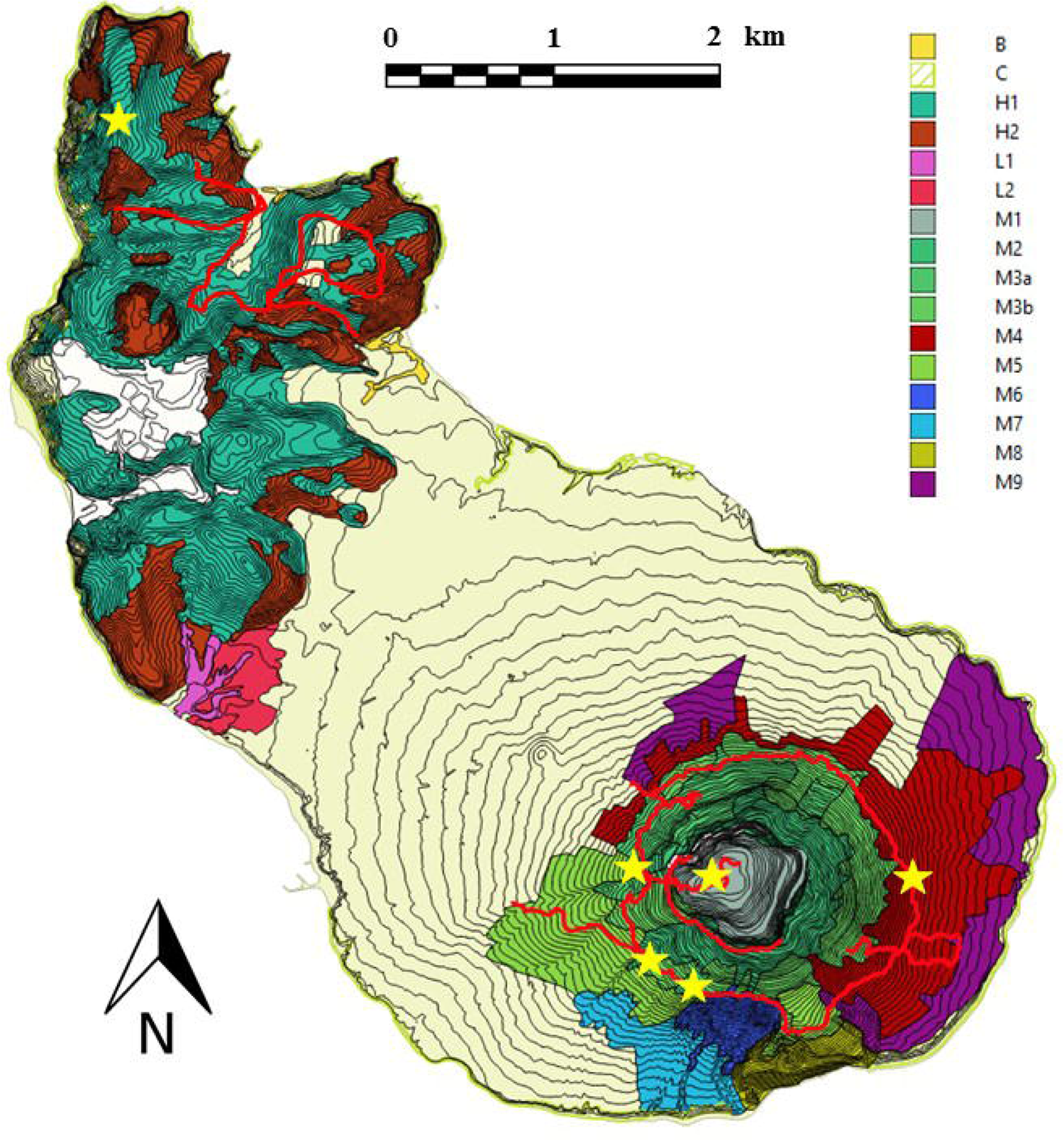
Detection probability (%) of *Alsophis rufiventris* in relation to sampling effort (number of surveys) on St. Eustatius based on transect surveys conducted in 2018 and 2019.

**Figure 7.**
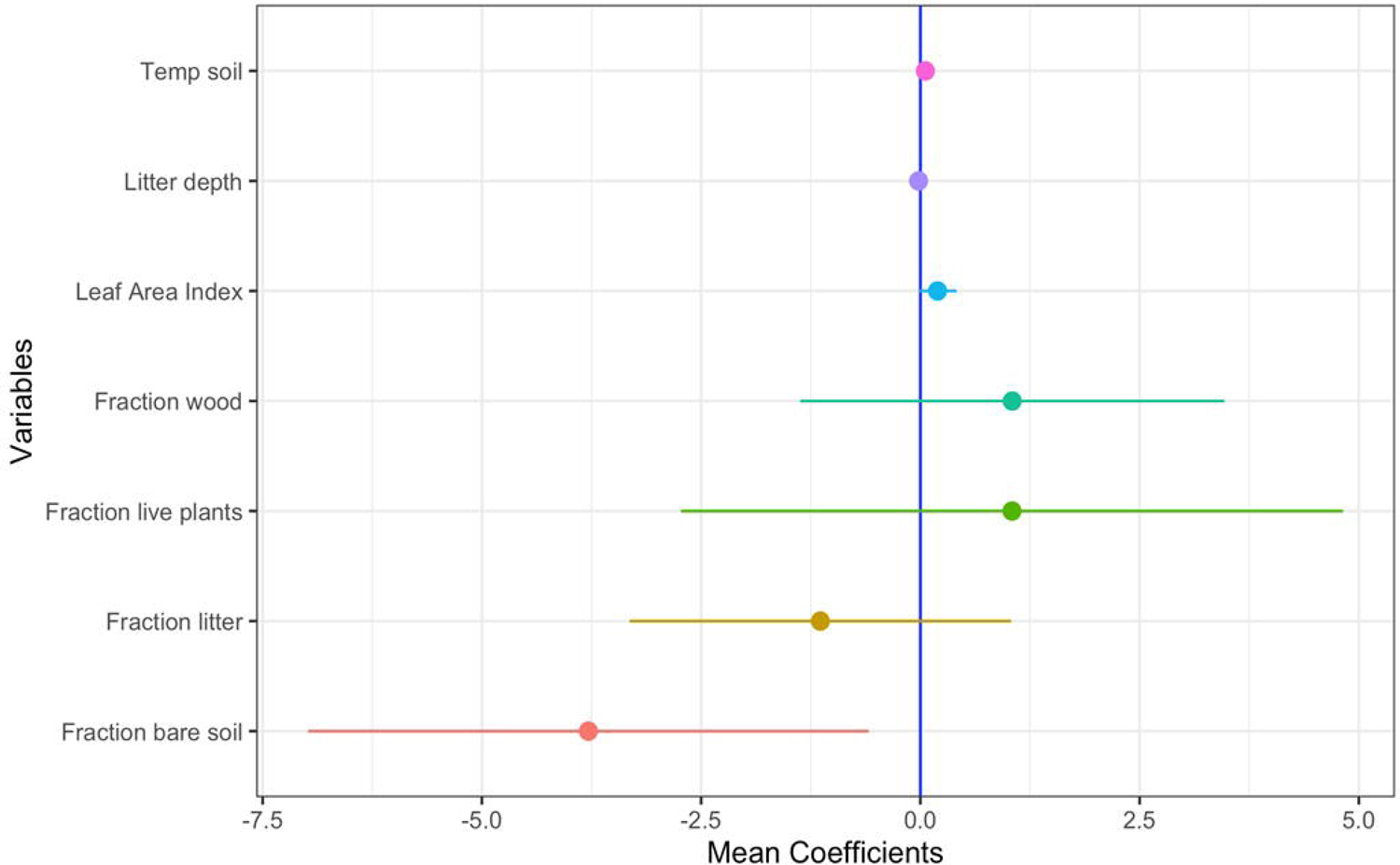
The effect of elevation on abundance (*λ*)estimates of *Alsophis rufiventris* on St. Eustatius based on transect surveys conducted in2018 and 2019.

**Figure 8.**
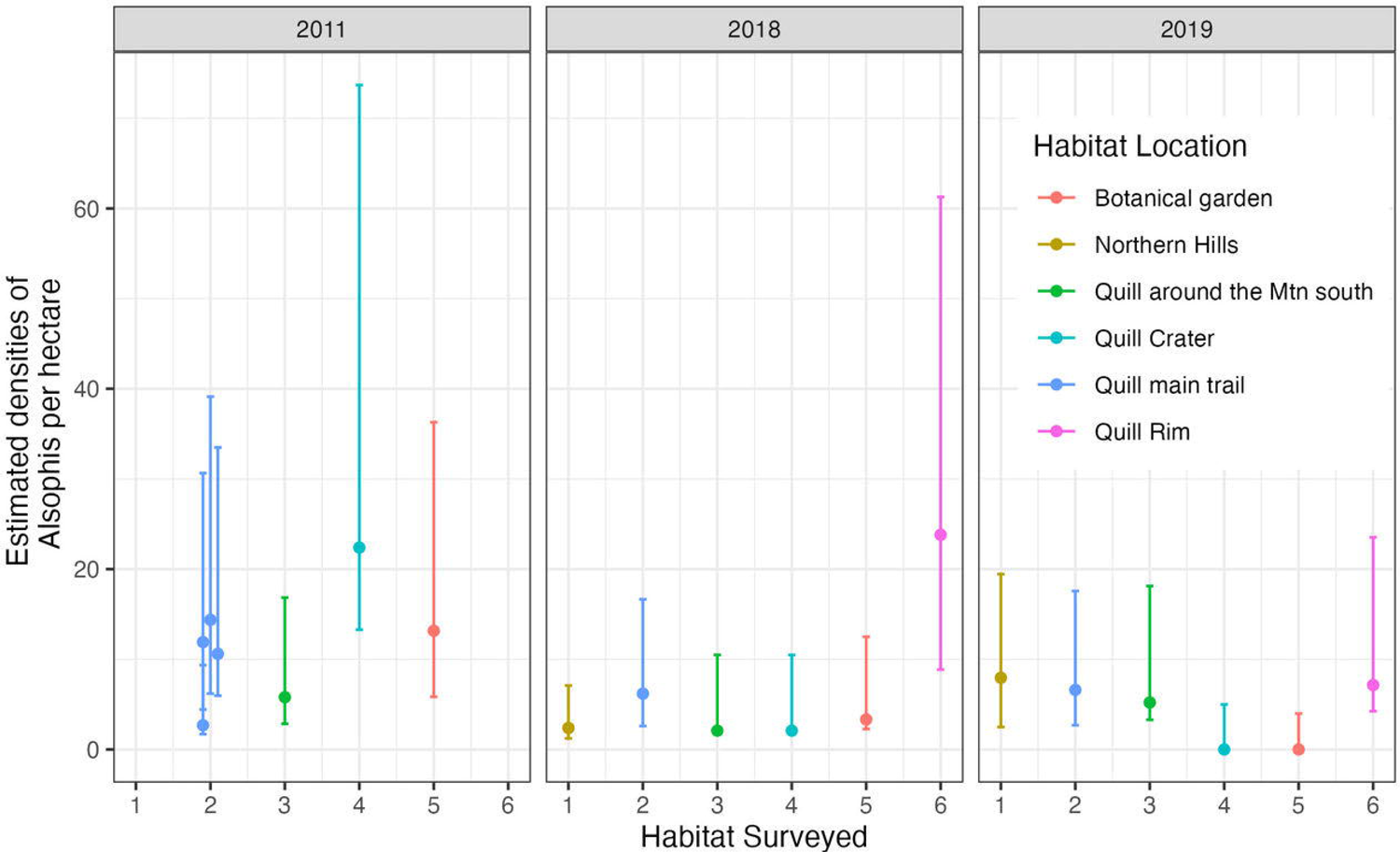
The effect of elevation on site occupancy estimates (ψ) of *Alsophis rufiventris* on St. Eustatius based on transect surveys conducted in 2018 (blue line) and 2019 (red line). Package: wiqid.

Habitat type did not appear to influence abundance (λ) and site occupancy (ψ) estimates in 2018 and 2019; note that the 95% confidence intervals for each habitat type are very wide and encompass each other. This was independent if the analysis was performed with ‘wiqid’ (Fig. 9a) or ‘unmarked’ software (Fig. 9b). Inside the crater, ψ estimates decreased from 0.98 (0.21 - 1.00 CI) in 2018 to 0.52 (0.21 - 0.82 CI) in 2019. λ estimates (average number of individuals per transect) were lowest in the northern hills (0.26 ± SD 0.17) and highest along the crater rim (2.87 ± 0.36) in 2018. λ estimates were lowest in the botanical garden/lower Quill slopes (3.25 ± SD 1.14) and highest along the main Quill trail (14.05 ± 5.40) in 2019. Occupancy rates differed among years, and the effects of covariates on detection estimates were only significant in 2018: survey time positively influenced detection probability (1.90 ± 0.26), whereas the effects of week of the survey (−0.20 ± 0.14), rain (−0.18 ± 0.18) and temperature (−0.08 ± 0.15) were negative. The effects of covariates on detection and occupancy estimates in 2019 were negligible (ΔAICc <2). Covariate models only shared survey time, week of survey and rainfall, the other variables only showed up in one of the models.

**Figure 9a.**
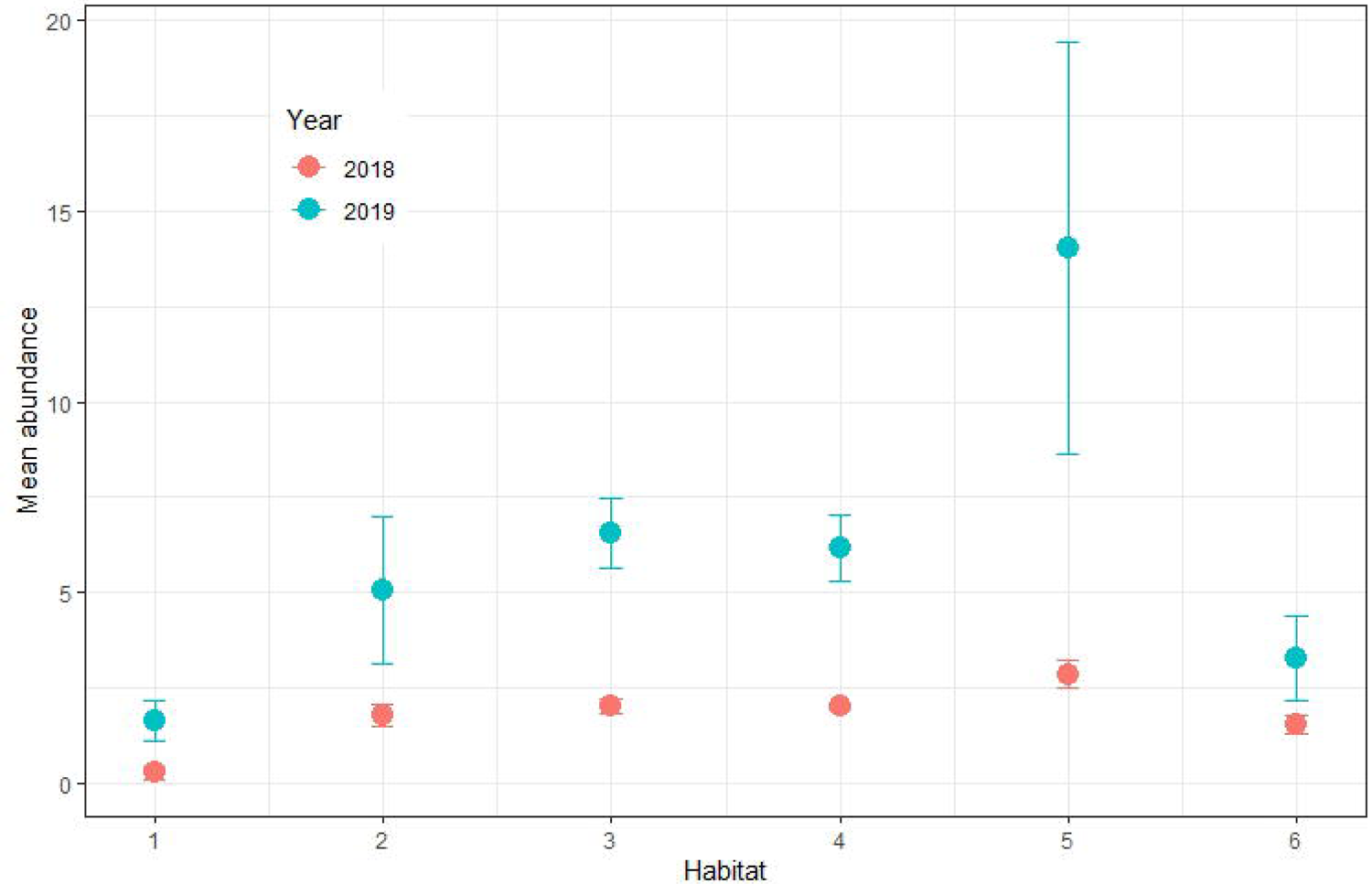
Site occupancy estimates per habitat of *Alsophis rufiventris* on St. Eustatius based on surveys conducted in 2018 and 2019. 1 = Northern hills; 2 = Quill main trail; 3 = Quill Around the Mountain S (upper); 4 = Quill crater; 5 = Quill rim; 6 = Botanical Garden/Around the Mountain S (lower). Package: wiqid.

**Figure 9b.**
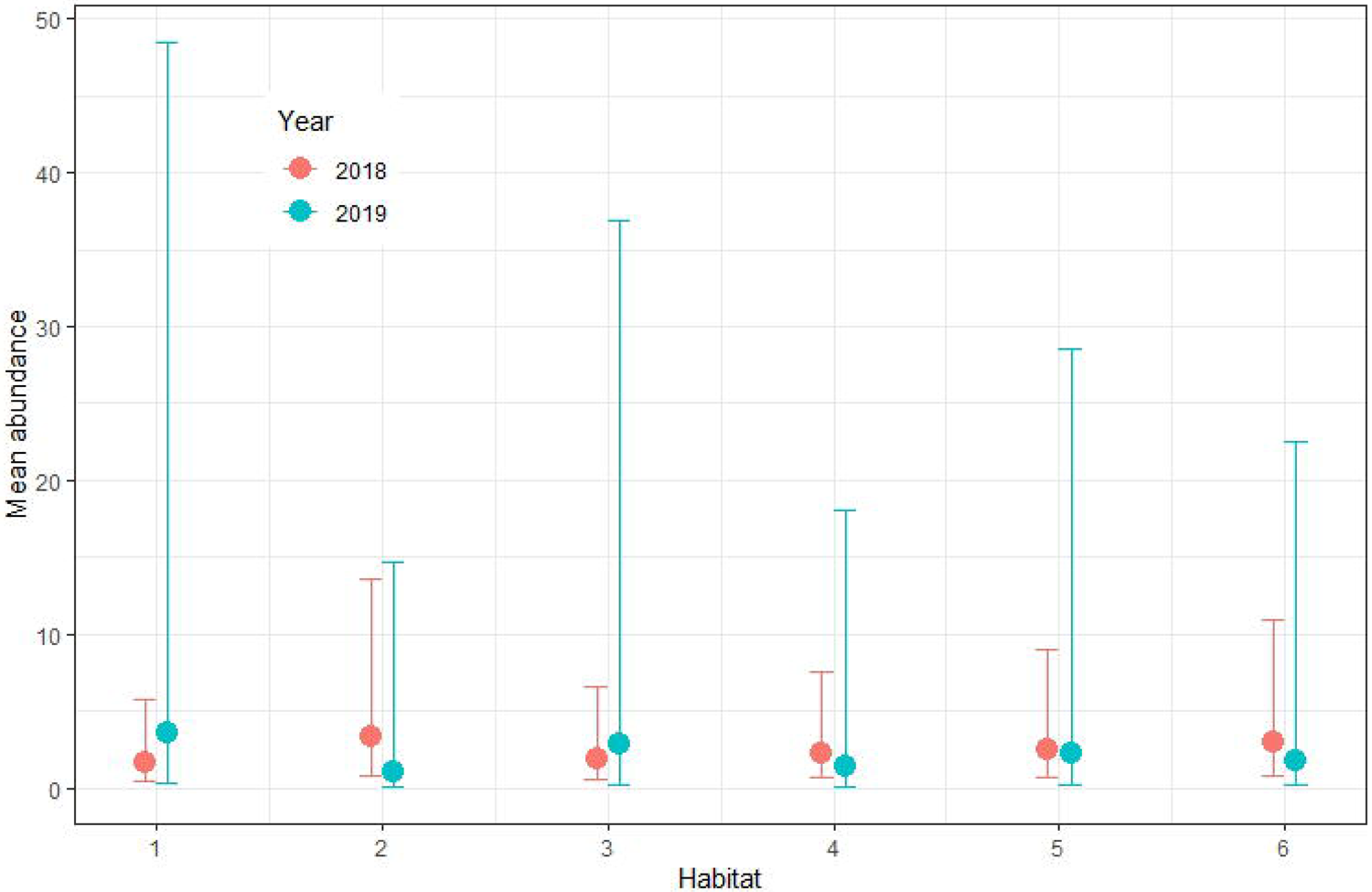
Site occupancy estimates per habitat of *Alsophis rufiventris* on St. Eustatius based on surveys conducted in 2018 and 2019. 1 = Northern hills; 2 = Quill main trail; 3 = Quill Around the Mountain S (upper); 4 = Quill crater; 5 = Quill rim; 6 = Botanical Garden/Around the Mountain S (lower). Package: unmarked.

## DISCUSSION

Quantitative estimates of abundance are essential for the recognition, management, and recovery of vulnerable, threatened, and endangered populations and species. Unfortunately, obtaining such estimates is often difficult, especially for rare organisms (Conroy et al., 2008). Worldwide, >82% of snake species have not been evaluated by the International Union for Conservation of Nature (IUCN) or are classified as data deficient (IUCN, 2020). Snakes are generally secretive animals with low abundances, leading to difficulty in detection and the perception that many species are impossible to monitor (Urbina-Cardona, 2008; Luja et al., 2008, Durso et al., 2011). Unfortunately, this has led to a lack of information about the status of most snake populations (Durso et al., 2011). We aimed to address this by producing the first quantitative assessment of hurricane impacts on an endangered snake species with an extremely restricted range.

Hurricanes can significantly alter the structure of biotic assemblages within hours of impact (Wiley and Wunderle, 1993; Widmer et al., 2004; Vilella and Fogarty, 2005; Schriever et al., 2009; Nicoletto 2013), and could cause local extirpation of some species (Schoener et al., 2017). The intensity and frequency of hurricanes has increased over recent decades (Emanuel, 2013, Trenberth 2005, Webster et al. 2005), and assessing their influence on biodiversity, although challenging, is of increasing importance. There is a general dearth of information on the effects of hurricanes on snake species, although Marroquín-Páramo et al. (2021) recorded a significant decrease in snake abundance and species richness in the disturbed tropical dry forest of Jalisco, Mexico, following hurricanes Jova (2011) and Patricia (2015). A decrease in predatory reptile species such as *A. rufiventris* could affect the local food web (Spiller and Schoener, 1998; Schoener et al., 2001).

Various studies have revealed differing responses of herpetofauna following hurricanes, with some species or assemblages increasing or decreasing their abundance, richness and diversity, while others remain unchanged (Reagan, 1991; Woolbright, 1991; Schoener et al., 2001; Gunzburger et al., 2010; Nicoletto 2013; Suazo-Ortuño et al. 2018b). The cumulative effects of two or more hurricanes usually result in a significant abundance decrease for specialist species, but an increase for generalist species (McCoid, 1996; Schriever et al., 2009; Selman, 2015). For example, in Louisiana following hurricanes Ivan (2004), Katrina and Rita (2005), *Thamnophis proximus* (Say in James, 1823) and *Agkistrodon piscivorus* (Lacépède, 1789) were observed to increase in abundance in levee and marsh habitats (Schriever et al., 2009). The structural effects of major hurricanes on forest ecosystems include tree damage/mortality (Eppinga and Pucko 2018) and an increase in leaf litter, which might affect the heterogeneity of the forest floor (Bellingham et al., 1996) and alter the structure and composition of herpetofauna assemblages, although this may be less pertinent on St. Eustatius given its small size and limited reptile diversity (Powell et al. 2005). Nevertheless, such changes may cause snakes to decline in abundance or disappear from some sites (Suazo-Ortuño et al., 2018a); indeed, our results show that post-hurricane racer density decreased dramatically inside the Quill crater compared to before the hurricanes. The racer population likely declined substantially within the first year post-hurricane, but exhibited signs of recovery after two years. This implies that the species may be resilient to hurricanes and could recover to pre-hurricane densities within approximately five years, provided no additional hurricanes make landfall on St. Eustatius during that time. Pre-hurricane data revealed that racers preferred sites without bare soil and with shallow litterfall depth. There was a small positive relationship between racer occurrence and soil temperature and fraction of green vegetation and wood litter on the ground, in addition to higher leaf area index. The higher leaf area index may suggest that racers are less likely to be observed in direct sunlight and prefer some amount of shade, however evaluating this variable independently showed no such pattern (GLM, binomial response, p >0.05).

Mean racer abundance increased across all habitats between 2018 and 2019, exhibiting similar patterns per habitat in both years. Specifically, abundance was highest in upper elevations of the Quill, especially along the crater rim, compared to Boven and the Botanical Garden/lower Quill slopes. This may be linked to greater quantities of prey such as *Anolis* lizards at higher elevations (Diaz et al., 2005). Alternatively, racers may prefer moist, unfragmented habitats at higher elevations with fewer shrubs and taller trees (Savit et al., 2005). Abundance estimates increased in line with increasing elevation in both years. We posit that the Quill’s higher elevational habitats (<600 m) are preferred by racers compared to the northern hills (296 m; DCNA 2014). This is also consistent with Daltry et al. (1997) who observed more racers in the Quill than the drier northern hills, and with other researchers who have selected the Quill to research racers (e.g., Savit et al., 2005; Zobel, 2016). Generally, however, racers are habitat generalists and have been observed across the entire island (pers. obs. HM). Our results also suggest that racers are more likely to be detected on cooler days, supporting our suggestion that the higher elevations of the Quill offer preferential habitat for the species. The effects of temperature on snake populations have been studied in temperate (Weatherhead et al., 2002) and Mediterranean climates (Zamora-Camacho et al., 2010), but we were unable to find relevant comparative studies of diurnal terrestrial snake species from the tropics. Moreover, it is possible that racer abundance and occupancy is influenced by other factors that were not accounted for during data collection (i.e., canopy cover, rock cover, leaf litter cover, prey abundance).

The Quill and Boven are separated by an airport and areas of agriculture and human habitation. Given that the parks likely support small, fragmented racer populations, if local extinction occurs in one area, natural recolonization may be inhibited by barriers of unsuitable habitat (Sewell et al., 2012). Additionally, we were unable to quantify the effects of invasive species which may impact *Alsophis* populations, however we note that free-roaming goats are present in both survey areas, with higher densities in the northern hills than the Quill (Madden, 2020). McCauley et al. (2006) suggest that snakes could be affected by free-roaming herbivores through trampling or loss of vegetation through grazing; the latter is certainly evident on the outer slopes of the Quill and the remaining forest fragments of Boven. A roaming animal control project is currently being initiated by the government of St. Eustatius (BES Reporter, 2020), part of which involves the installation of a fence across the southern boundary of Boven National Park followed by the removal of non-native herbivores. This should result in the recovery of vegetation and reduce the risk of disturbance/trampling of racers by roaming goats. Black rats are also present in various densities across all vegetation types on St. Eustatius, but are more prevalent in forested areas (Madden et al., 2020). Thus the Quill is an ideal habitat for black rats as well as racers, and the coexistence of both species could have negative impacts on the racer population, as suggested by Daltry et al. (1997). Currently there are no plans to control or eradicate rats or free-roaming livestock from the Quill, which would be effective short-term conservation measures for the persistence of the racer population. Herpetofauna perform various regulating ecological functions in forest ecosystems (Cortés-Gomez et al., 2015) and can modify forest processes such as regeneration and nutrient cycling (Felix et al., 2004). Secondary dry tropical forest ecosystems play a buffering role that promote herpetofauna resilience, thus their importance for racer persistence cannot be underestimated, especially in line with increasing hurricane frequency and intensity (Marroquín-Páramo et al. 2021). Long-term monitoring to assess hurricane impacts, as well as changes in abundance and occupancy, will play an important role in the conservation of racers within St. Eustatius’ forest ecosystems, especially under the predicted future scenarios of anthropogenic climate change.

## ACKNOWLEDGEMENTS

We are grateful to St. Eustatius National Parks Foundation for allowing us to conduct surveys within the parks and for granting permission to handle and PIT tag snakes in 2019. We thank CNSI for logistical support. We acknowledge the support of Tim van Wagensveld during 2018 and 2019 internships, and thank RAVON for the internship opportunity. We also thank Frank Rivera-Milán for his advice during 2018 surveys. We are grateful to F. Toledo-Rodríguez, Hernández-Velázquez, F.I., E.G. Martínez-Cebollero, E.M. Pérez-Rivera, S.M. Davila-Vázquez, S.Y. Lindsay-Zenón, J.R. Rodríguez-Reyes, K. Rodríguez-Rodríguez, and C. Figuerola for participating in 2011 fieldwork. Fieldwork in 2011 was funded by a National Science Foundation grant: PHRD 0734826. We greatly appreciate the support of staff and volunteers from STENAPA: Kate Walker, Sapphira Looker and Randy Stotesbery. We thank Wouter Koeneman and Dominique Koning from the Old Gin House for the superb lodging.

## Notes

### Competing Interest Statement

The authors have declared no competing interest.

