## supplementary tables for "Find me if you can: Pre- and Post-hurricane Densities of the Red-bellied Racer (*Alsophis rufiventris*) on St. Eustatius, and a review of the genus in the Caribbean"

**TITLE**

**Journal**: South American Journal of Herpetology

**Table S1.** Coordinates and elevation of areas surveyed for *Alsophis rufiventris* on St. Eustatius in 2011, 2018, and 2019.

| **Site Name** | **Pre- post Hurricane** | **Year of survey** | **North Coordinate** | **West Coordinate** | **Start Elevation** | **Length (m) and area surveyed** | **Vegetation type** |
| --- | --- | --- | --- | --- | --- | --- | --- |
| Quill main trail: *Pisonia-Eugenia* mountains | Pre | 2011 | 17°28’38.6” N | -062°58’12.1” W | 273 | 372m:  18600m^2^ | M5 |
| Quill main trail: *Pisonia-Eugenia* mountains | Pre | 2011 | 17°28’35.3” N | -062°58’11.0” W | 277 | 487m:  9740m^2^ | M5 |
| Quill southern trail: *Pisonia-Eugenia, Chionanthus-Nectandra* and *Capparis-Antirhea* mountains | Pre | 2011 | 17°28’19.9” N | -062°57’56.5” W | 314 | 345m:  17250m^2^ | M6 |
| Quill main trail: *Pisonia-Eugenia* mountains | Pre | 2011 | 17°28’22.8” N | -062°58’30.0” W | 269 | 330m:  6600m^2^ | M5 |
| Quill crater and inner slopes: *Myrcia-Quararibea* | Pre | 2011 | 17°28’39.3” N | -062°57’54.4” W | 317 | 336m:  16800m^2^ | M1 |
| Quill main trail: *Pisonia-Eugenia* mountains | Pre | 2011 | 17°28’40.8” N | -062°58’47.2” W | 358 | 134m:  2680m^2^ | M5 |
| Botanical Garden, Quill lower SW flanks: *Capparis-Pisonia* mountains | Pre | 2011 | 17°28’44.4” N | -062°57’15.9” W | 192 | 494m:  9880m^2^ | M4 |
| Northern Hills: *Pisonia-Justicia* and *Pisonia-Bothriochloa* hills | Pre | 2011 | 17°31’16.7” N | -062°59’47.8” W | 254 | occasional | H1,H2 |
| Northern Hills: *Pisonia-Justicia* and *Pisonia-Bothriochloa* hills | Post | 2018, 2019 | 17°30’41.7” N | -062°59’07.4” W | 13 | 4000m: 48000m^2^ | H1, H2 |
| Quill main trail: *Pisonia-Eugenia* mountains | Post | 2018, 2019 | 17°28'34.9" N | -062°58'28.5" W | 155 | 2020m: 24240m^2^ | M5 |
| Quill southern trail: *Pisonia-Eugenia, Chionanthus-Nectandra* and *Capparis-Antirhea* mountains | Post | 2018, 2019 | 17°28'24.2" N | -062°58'04.4" W | 275 | 800m: 9600m^2^ | M5, M3b, M6 |
| Quill crater and inner slopes: *Myrcia-Quararibea* | Post | 2018, 2019 | 17°28'41.9" N | -062°57'50.7" W | 296 | 800m: 9600m^2^ | M1 |
| Quill rim: *Coccoloba-Chionanthus* and *Chionanthus-Nectandra* mountains | Post | 2018, 2019 | 17°28'37.8480" N | -062°58'01.2” W | 393 | 900m: 10800m^2^ | M2, M3a |
| Botanical Garden, Quill lower SW flanks: *Capparis-Pisonia* mountains | Post | 2018, 2019 | 17°28'28.8” N | -062°57'09.0” W | 80 | 1000m: 12000m^2^ | M4 |

**Table S2.** *Alsophis rufiventris* marked and recaptured individuals with PIT tags during surveys on St. Eustatius in 2018 and 2019. Red text indicates recapture.

| **PIT tag ID** | **Date PIT-tagged** | **Time** | **Sex** | **Total length (cm)** | **Weight (g)** | **Recapture Y/N (date)** |
| --- | --- | --- | --- | --- | --- | --- |
| 6862 | 10/3/2016 | 08:29 | F | 84.9 | - | N |
| *6862* | *10/3/2016* | *10:24* | *F* | *85.0* | *160* | *2/5/2018* |
| 5012 | 4/1/2019 | 10:49 | M | 93.8 | 125 | N |
| 5039 | 4/4/2019 | 11:17 | M | 85.0 | 130 | N |
| 5033 | 4/4/2019 | 12:15 | M | 81.6 | 140 | N |
| 6877 | 4/9/2019 | 09:13 | M | 64.1 | 95 | N |
| 6890 | 4/16/2019 | 09:00 | F | 82.2 | 155 | N |
| 6827 | 4/16/2019 | 17:56 | M | 77.4 | 120 | N |
| 6878 | 4/18/2019 | 09:39 | F | 107.0 | 155 | N |
| 6883 | 4/18/2019 | 15:32 | M | 104.6 | 170 | N |
| 6884 | 4/26/2019 | 17:44 | M | 92.5 | 175 | N |
| 6827 | 5/23/2019 | 10:29 | M | 85.3 | 120 | N |
| 3819 | 5/23/2019 | 16:15 | M | 89.4 | 145 | N |
| *3819* | *5/23/2019* | *16:32* | *M* | *90.1* | *145* | *6/11/2019* |
| 3843 | 5/23/2019 | 16:57 | F | 85.5 | 155 | N |
| 3804 | 5/30/2019 | 15:41 | M | 82.9 | 100 | N |
| 3757 | 6/3/2019 | 12:10 | M | 89.2 | 170 | N |
| 3842 | 6/4/2019 | 12:50 | M | 88.6 | 130 | N |
| 3794 | 6/4/2019 | 13:29 | M | 88.4 | 110 | N |
| 3772 | 6/4/2019 | 14:10 | F | 89.5 | 115 | N |
| 3811 | 6/4/2019 | 14:44 | M | 88.8 | 135 | N |
| 3750 | 6/5/2019 | 15:28 | M | 88.0 | 110 | N |
| 3825 | 6/5/2019 | 17:21 | M | 77.0 | 95 | N |
| 3805 | 6/7/2019 | 10:09 | M | 83.8 | 80 | N |

**Table S3.** List of authorship and year of publication of all taxon names mentioned in the text.

Barbour, T. 1914. *Eleutherodactylus johnstonei*. In Species 2000 & ITIS Catalogue of Life: 2019, Catalogue of Life.

Barbour, T. 1915. Recent notes regarding West Indian reptiles and amphibians. *Proceedings of the Biological Society of Washington* 28: 71–78.

Cope, E.D. 1869. Seventh contribution to the herpetology of tropical America. *Proceedings of the American Philosophical Society* 11(81): 147–192.

Cope, E.D. 1879. Eleventh contribution to the herpetology of tropical America. *Proceedings of the American Philosophical Society* 18: 261–277.

Daudin, F. 1802. Histoire Naturelle, Générale et Particulière des Reptiles, Vol. 4. F. Dufart, Paris.

Daudin, F. 1803. *Indotyphlops braminus*. In Species 2000 & ITIS Catalogue of Life: 2019, Catalogue of Life.

Duméril, A., Bibron, G. & Duméril, A. 1854. *Alsophis rufiventris*. Published in: Garman, S. 1887. On West Indian reptiles in the Museum of Comparative Zoology at Cambridge, Mass. *Proceedings of the American Philosophical Society* 24: 278–286.

Geoffroy Saint-Hilaire, É. 1818. *Herpestes javanicus.* In Species 2000 & ITIS Catalogue of Life: 2019, Catalogue of Life.

Kuhl, H. 1824. Sur les reptiles de Java. *Bulletin de la Société Sciences Nat*, Paris 2:79–83.

Laurenti, J.N. 1768. Specimen medicum, exhibens synopsin reptilium emendatam cum experimentis circa venena et antidota reptilium austracorum, quod authoritate et consensu. Vienna, Joan. Thomae, 217 pp.

Lacépède, Comte de. 1789. Histoire naturelle des serpens. 1–144 + 1–527 p. Paris.

Lazell, J.D. 1972. The anoles (Sauria: Iguanidae) of the Lesser Antilles. Bulletin of the Museum of Comparative Zoology. Harvard 143 (1): 1–115.

Linnaeus, C. 1758. Systema naturae per regna tria naturae, secundum classes, ordines, genera, species, cum characteribus, differential, synonymis, locis, Tomus I. Editio decima, reformata. Laurentiis Salvii, Holmiae. DOI

Reinhardt, J.T. 1843. Beskrivelse af nogle nye Slangearter. Det Kongelige Danske Videnskabernes Selskabs Naturvidenskabelige og Mathematiske Afhandlinger, 10, 233–279.

Reinhardt, J.T., Lütken, P. 1862. *Borikenophis portoricensis.* In Species 2000 & ITIS Catalogue of Life: 2019, Catalogue of Life.

Say, T. 1823. *Coluber obsoletus*. In E. James, ed. Account of an expedition from Pittsburgh to the Rocky Mountains, performed in the years 1819 and ’20. Logman, Hurst, Rees, Ovme and Brown, London. P. 140

Schlegel, H. 1837. Essai sur la physionomie des serpens. Partie Générale: xxviii 251 S. Partie Descriptive: 606.

Thomas, R. 1966. Leeward Islands Typhlops (Reptilia, Serpentes). *Proceedings of the Biological Society of Washington* 79: 255–265.
